# Correlated variability in primate superior colliculus depends on functional class

**DOI:** 10.1101/2021.09.15.460545

**Authors:** Leor N. Katz, Gongchen Yu, James P. Herman, Richard J. Krauzlis

## Abstract

Correlated variability (spike count correlations, r_SC_) in a population of neurons can constrain how information is read out, depending on behavioral task and neuronal tuning. Here we tested whether r_SC_ also depends on neuronal functional class. We recorded from populations of neurons in macaque superior colliculus (SC), a structure that contains well-defined functional classes. We found that during a guided saccade task, different classes of neurons exhibited differing degrees of r_SC_. “Delay class” neurons displayed the highest r_SC_, especially during the delay epoch of saccade tasks that relied on working memory. This was only present among Delay class neurons within the same hemisphere. The dependence of r_SC_ on functional class indicates that subpopulations of SC neurons occupy distinct circuit niches with distinct inputs. Such subpopulations should be accounted for differentially when attempting to model or infer population coding principles in the SC, or elsewhere in the primate brain.

## Introduction

Neuronal responses to similar sensorimotor settings are variable, and often correlated between neurons of a population. Such correlated variability (spike count correlations, r_SC_) is an important topic in systems neuroscience because the degree of r_SC_ can have profound implications on population codes (Abbott and Dayan, 1999; Shadlen and Newsome, 1998; Sompolinsky et al., 2001; Zohary et al., 1994). When making judgements about sensory events, perceptual accuracy can be dramatically impaired if the shared variability in neuronal responses mimics the stimulus-driven activity (Averbeck et al., 2006; Cohen and Kohn, 2011; Moreno-Bote et al., 2014). Identifying the factors that determine r_SC_ during behavioral tasks is therefore critical for understanding how neurons interact to guide behavior (Kohn et al., 2016; Nienborg et al., 2012).

What are the factors that determine r_SC_? Mechanistically, the presence of r_SC_ in a population is evidence of common inputs onto the neurons. The origin of such inputs, however, is still debated (Bondy et al., 2018; Kanitscheider et al., 2015; Shadlen and Newsome, 1998). Seminal experiments (Britten et al., 1992; Zohary et al., 1994) had attributed the presence of r_SC_ to shared noise in the sensory inputs, which led to the term “noise correlation” in reference to r_SC_ (Averbeck et al., 2006; Cohen and Kohn, 2011; Shadlen and Newsome, 1998). Today, however, the term “noise correlations” is largely appreciated as a misnomer as the shared variability is driven less by noise on sensory inputs and instead, by signals that had yet to be identified at the time. For example, r_SC_ can be driven by global fluctuations in population activity that are not noise (Ecker et al., 2014; Goris et al., 2014; Harris and Thiele, 2011; Schölvinck et al., 2015), and by task-related feedback signals that target specific neuronal populations (Bondy et al., 2018; Cohen and Newsome, 2008; Cumming and Nienborg, 2016). Consistent with the idea that spike-count correlations are largely due to feedback signals, changes in r_SC_ have been observed following a variety of experimental manipulations aimed at influencing cognitive states such as attention (Cohen and Maunsell, 2009; Mitchell et al., 2009), perceptual learning (Gu et al., 2011; Ni et al., 2018), decision-making (Bondy et al., 2018; Nienborg et al., 2012) and task context (Bondy et al., 2018; Cohen and Maunsell, 2009).

Given the specific anatomical routes by which task-related feedback is applied within a brain region versus feedforward inputs, it might be expected that individual neurons would exhibit different degrees of r_SC_ that are related to their function, processing stage or association with either feedforward or feedback projections. We tested this hypothesis in the primate superior colliculus (SC) because SC neurons are readily classified into distinct functional classes based on their spiking response properties during saccade tasks. Neurons defined as either visual, visual-movement, movement, or delay correspond to different locations within the SC circuitry and are linked to distinct sources of input and output (Krauzlis et al., 2013; Basso and May, 2017; Gandhi and Katnani, 2011; Wurtz and Albano, 1980). Distinct sources of input may carry signals that differ in their degree of correlation as well as their dependence on task conditions, and may give rise to distinct levels of r_SC_ across the different classes of SC neurons.

We recorded from populations of SC neurons and compared the degree of r_SC_ within and between functional classes over multiple epochs of an oculomotor working memory task. We found that neuron pairs classified by function exhibited different degrees of r_SC_. Moreover, neurons belonging to the “Delay” class displayed particularly high correlation values during saccade tasks that relied on working memory. Such variability in r_SC_ indicates that sub-populations of neurons occupy distinct niches within the SC network, and likely receive distinct modulatory inputs from upstream regions. We conclude that the circuit niche of subpopulations of neurons, at least as reflected by their functional class, is important to consider to accurately inform population coding models in SC, or elsewhere in the brain.

## Results

We recorded from the superior colliculus (SC) of two nonhuman primates (*Macaca Mulatta*) trained to perform saccadic eye movements. During each recording session, monkeys performed visually guided and memory-guided saccades on separate, randomly interleaved trials (Figure 1B). Whereas the saccade target remained visible during the delay epoch of the visually guided saccade condition, it was absent from the memory-guided version, requiring the monkey to rely on their working memory of target location to guide the saccade. Saccade target location on each trial was selected randomly from among multiple locations, including a location within the envelope of neuronal receptive fields (RF), termed the “in-RF” target.

**Figure 1:**
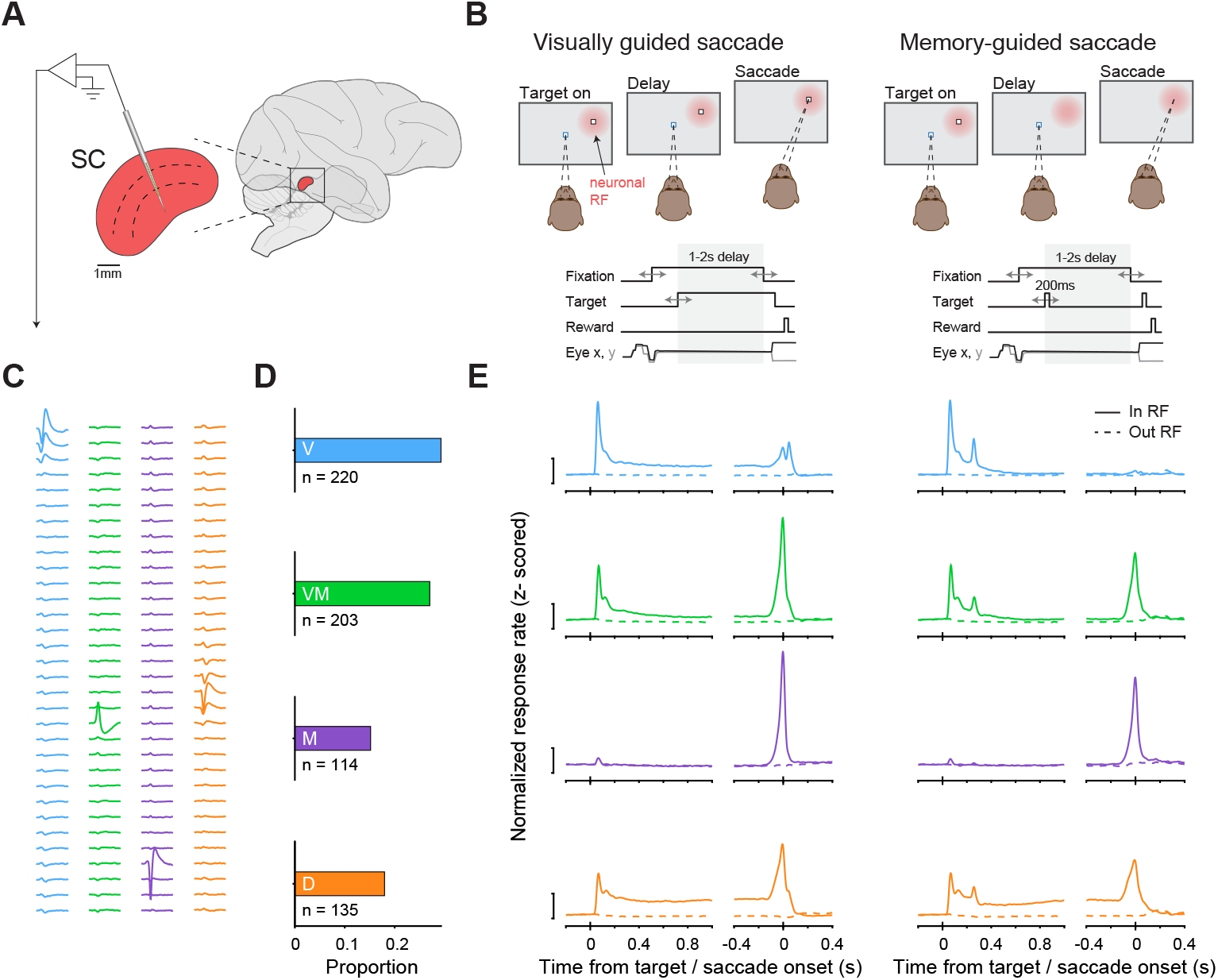
SC neurons were readily classified into functional classes during a guided saccade task. **(A)** A multi-channel probe was positioned to primarily target the intermediate layers of SC in macaques performing a guided saccade task. **(B)** Monkeys performed either visually guided (left) or memory-guided saccades (right). Top: in either task condition, the subject was required to maintain fixation during both target presentation and a 1 – 2 s delay period. The offset of the fixation point cued the monkey to saccade to the target to obtain a juice reward. The key difference between task conditions is that the target did not remain visible during the delay period of memory-guided saccade trials. Visually guided and memory-guided saccade trials were interleaved, and the target could appear within the envelope of neuronal receptive fields (RF, indicated by the red patch) or elsewhere. Bottom: timing of task events. **(C)** Example waveforms on each probe channel (rows) from isolated neurons from a single recording session, one set from each functional class (colors matched to panels D & E). **(D)** Proportion of neurons recorded from each functional class: Visual (V); Visual-Movement (VM); Movement (M); and Delay (D). **(E)** Population average of z-score normalized responses for each of the classes for saccades into the neuronal RF (“in RF”) and out (“out RF”), relative to target onset and saccade onset, during visually guided (left) and memory-guided saccade trials (right).

A multichannel recording probe was positioned to span multiple layers of the SC such that populations of neurons with different functional properties were recorded simultaneously (Figure 1A, C). Overall, we recorded from 751 neurons with well-defined RFs over 45 recording sessions (Monkey R: 309 neurons over 16 sessions; Monkey P: 442 neurons over 29 sessions). Results presented below were quantitatively consistent across the two monkeys and were therefore combined.

### SC neurons were readily classified into functional classes

SC neurons were classified into distinct functional classes following well-established criteria, based on their responses during the visual, delay, and movement periods of the memory-guided saccade task (see Methods). Applying statistical criteria used previously (McPeek and Keller, 2002), we assigned each neuron to a functional class: “Visual” (29%); “Visual-Movement” (27%); “Movement” (15%); and neurons exhibiting persistent activity during the delay epoch of the memory-guided saccade task, termed “Delay” neurons (18%) (Figure 1D). The mean response of each sub-population was consistent with previous reports (Basso and Wurtz, 1998; Herman and Krauzlis, 2017; McPeek and Keller, 2002) for either visually or memory-guided saccades trials (Figure 1E).

### Correlated variability in SC depended on functional class

The classification of SC neurons into sub-populations enabled us to test whether different classes of SC neurons exhibited different degrees of correlated variability. For neurons within a hemisphere, r_SC_ was only computed for neuron pairs with overlapping RFs, netting 5339 pairs of simultaneously recoded neurons. Within each functional class, our dataset included 627, 590, 144, and 280 pairs from the Visual, Visual-Movement, Movement, and Delay classes, respectively. Across functional classes, our dataset included an additional 3698 pairs.

As an initial characterization of correlated variability among SC neurons, we measured the time course of spike count correlations during both types of saccade tasks, computed over all SC neuron pairs regardless of functional class. The average r_SC_ during visually guided saccade trials (Figure 2A) was modest (approximately 0.06) with a brief increase in r_SC_ shortly after target onset (during the “visual epoch”, see figure panel), followed by a return to ∼0.06 during the “delay epoch”. A modest reduction in r_SC_ was noted during the “movement epoch”. Despite these variations, mean r_SC_ across pairs remained significantly above zero in every time bin (p < 0.001, Student’s t-test, corrected). The time course of r_SC_ during memory-guided saccades followed a similar pattern (Figure 2B).

**Figure 2:**
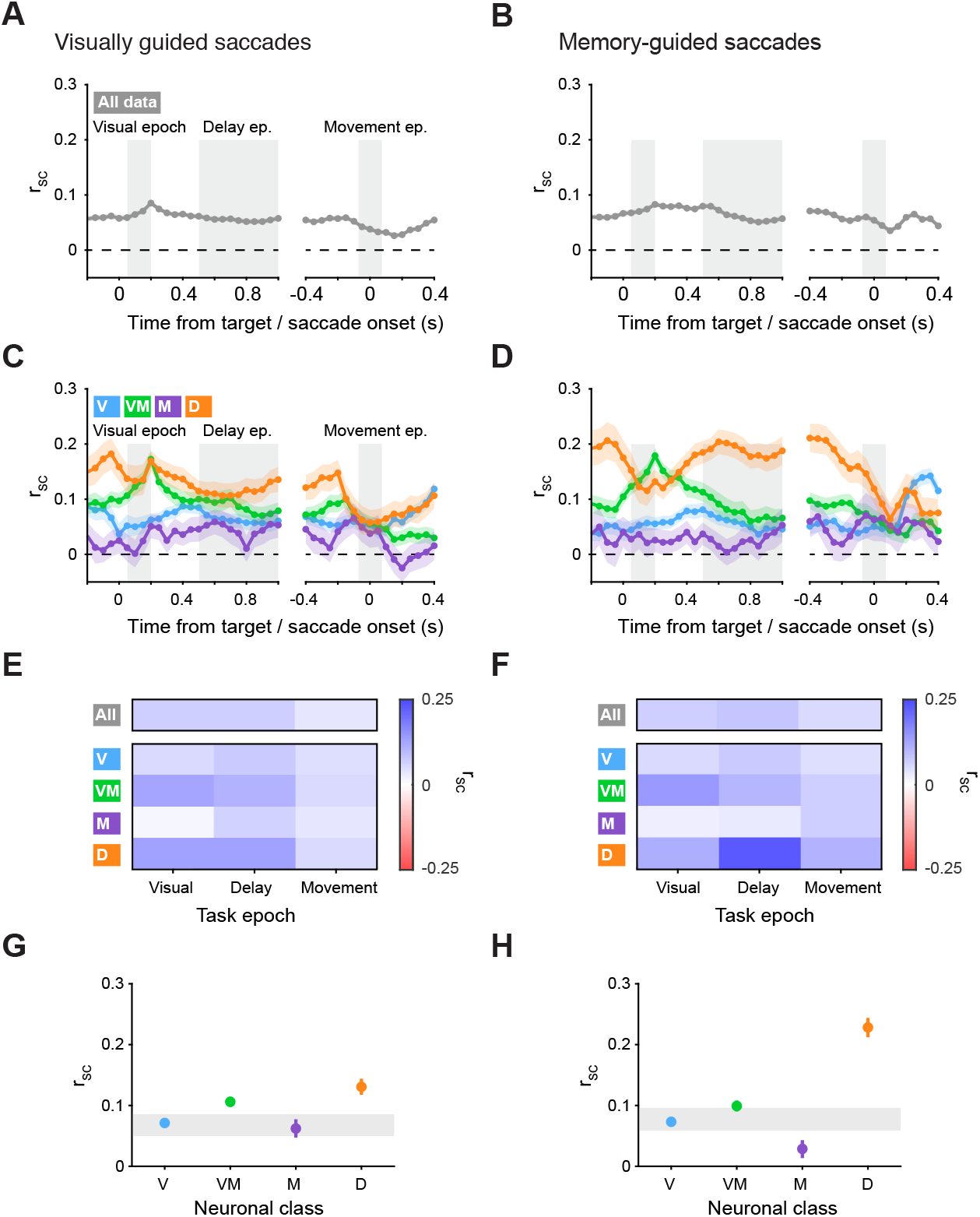
r_SC_ in SC depended on functional class. **(A-B)** Time-course of correlated variability (r_SC_) measured across all SC neurons collapsed across class during visually guided (A) and memory-guided (B) saccade trials, relative to target and saccade onset (150 ms bins in 50 ms steps). Three key epochs used for subsequent analysis are indicated: visual epoch (50-200ms after target onset); delay epoch (500-1000ms after target onset); and movement epoch (75ms before to 75ms after saccade onset). **(C-D)** Time-course of r_SC_ measured for distinct subpopulations of SC neuron pairs, classified as either Visual (V), Visual-Movement (VM), Movement (M), or Delay (D). Same format as A-B. **(E-F)** Top bar: Mean r_SC_ in each epoch for SC neuron pairs collapsed across class during visually guided (E) and memory-guided (F) saccade trials. Bottom matrix: Mean r_SC_ in each epoch for SC neuron pairs from each functional class. **(G-H)** Mean r_SC_ values for each functional class during the delay epoch of visually guided (G) and memory-guided (H) saccades. Error bars indicate 1 SEM, bootstrapped. Gray bar reflects 95% confidence intervals on the mean of r_SC_ values for pairs drawn from a class-blind “null” distribution, bootstrapped.

Partitioning SC neuron pairs into distinct functional classes revealed time courses of r_SC_ that varied substantially across neuronal class (Figure 2C, D), and were strikingly different from the r_SC_ computed on all SC pairs independent of class (Figure 2A, B). To statistically quantify how r_SC_ values varied across conditions we considered r_SC_ values in the three key task epochs indicated on Figure 2A-D: the visual epoch, the delay epoch, and the movement epoch. We found that for visually guided saccades (Figure 2E), the degree of r_SC_ depended strongly on task epoch, on neuronal class, and on the interaction between them (all p < 0.001, 2-way ANOVA). The highest values of r_SC_ were exhibited by neurons from the Visual-Movement and Delay classes, predominantly during the visual and delay epochs of the task. During the movement epoch, in contrast, neurons from neither class exhibited strong correlated activity. For memory-guided saccades (Figure 2F), a similar pattern was observed, with significant effects of task epoch, neuronal class, and the interaction between them (all p < 0.001, 2-way ANOVA). However, there was a conspicuous increase in r_SC_ for Delay class neurons during the delay epoch of the memory guided saccade task that was not observed during the visually guided task.

The dependence of r_SC_ on functional class was particularly striking during the delay epoch. Targeted analysis of this epoch revealed clear differences in r_SC_ across classes (Figure 2G, H). Mean r_SC_ values during visually guided saccade trials were positive in each neuronal class individually (Figure 2G) and significantly different from a matched trial-shuffled (null) distribution (Supplementary figure 1A). Moreover, r_SC_ values across classes significantly differed from one another (p < 0.001, ANOVA). Similar results were obtained for memory-guided saccade trials; neuron pairs in each of the four classes exhibited mean r_SC_ values that were significantly larger than zero (Figure 2H, Supplementary figure 1B) and significantly different from one another (p < 0.001, ANOVA), but with a much larger difference between pairs in the Delay class and in all other classes.

Thus, neurons that belong to distinct functional classes exhibited differing degrees of r_SC_, and these depended on task epoch and saccade condition. Spike count correlations were highest in Delay class neurons, and this was especially pronounced during the delay epoch of memory-guided saccades.

### Dependence of r_SC_ on functional class was not due to firing rate, distance, signal correlation or reaction time

Correlated activity is thought to arise through common input to a population of neurons. However, other factors may also affect the degree of measured r_SC._ Neuron pairs with higher firing rates tend to exhibit larger measured r_SC_, as do pairs that are physically closer (Bair et al., 2001; Cohen and Kohn, 2011; Kohn and Smith, 2005). To test whether these factors might account for our findings, we performed a mean-matching procedure (Churchland et al., 2010; Ruff and Cohen, 2014) to control for firing rate and distance between pairs (Methods). We found that even when these two factors were matched across neuronal classes, significant variation of r_SC_ across classes persisted (Supplementary Figure 2A). We further tested whether the variation in r_SC_ across classes could be accounted for by variation in relationship between signal correlations (r_Signal_) and r_SC_ (Cohen and Kohn, 2011; Kohn and Smith, 2005), and found no systematic variation corresponding to our measured r_SC_ across classes (Supplementary Figure 2B). Correlations between neuronal responses and saccadic reaction times did not explain our findings either, as these were not significantly different across classes (p > 0.05, Kruskal-Wallis). Thus, the differences in r_SC_ strength across neuronal classes were not explained by differences in firing rate, distance, r_Signal_ to r_SC_ relationship, or saccadic reaction times, and instead, points towards distinct sources of common input to the functional classes of SC neurons.

### r_SC_ within and between functional classes

Having established that pairs of SC neurons exhibit differing degrees of r_SC_ within each functional class, we next assessed the degree of pairwise correlations between classes. We quantified the degree of r_SC_ between neurons belonging to different functional classes, during the three epochs of the task (Figure 3). The between-class correlation values appear on the off-diagonal of the matrix (and consist of class-pairs: V-VM, V-M, V-D, VM-M, VM-D, and M-D, where V, VM, M and D stand for Visual, Visual-Movement, Movement and Delay, respectively). For reference, we also include the within-class correlation values (same data shown previously in Figure 2E and 2F); these appear on the diagonal (and consists of four pairs: V-V, VM-VM, M-M, and D-D).

**Figure 3:**
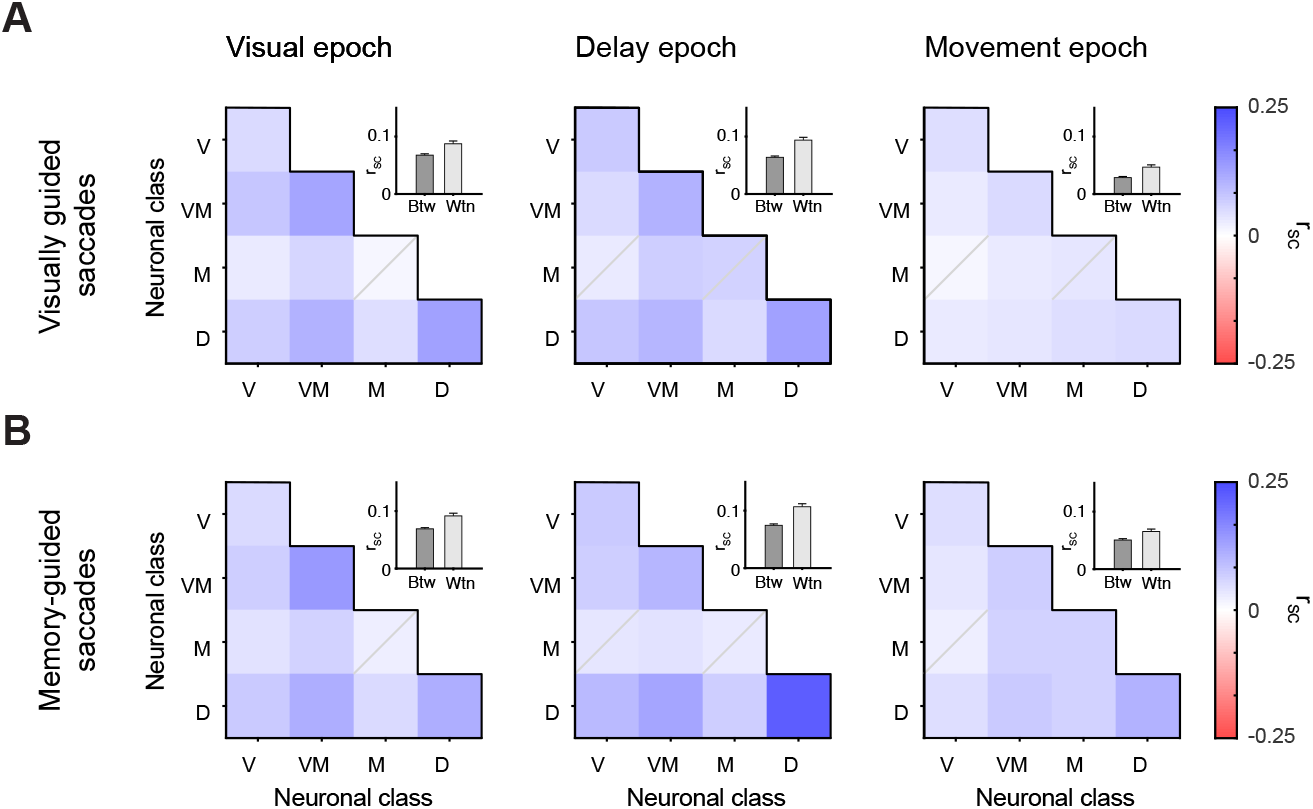
r_SC_ was larger within class than between classes. **(A-B)** Heatmaps of mean r_SC_ of neuron pairs between functional classes for visually guided (A) and memory-guided (B) saccades, one for each task epoch. Elements on the diagonal of the heatmap indicate within-class r_SC_ values, off diagonal elements indicate between-class r_SC_. Elements for which the mean r_SC_ of neuronal pairs was not statistically significantly different from their trials-shuffled (null) distribution are indicated by a gray diagonal on the element (p > 0.05, Student’s t-test, corrected). Insets show the mean r_SC_ across all within-class groups versus all between-class group for each epoch. Asterisks denote statistical significance: *** p < 0.001; ** p < 0.01; * p < 0.05.

Neurons exhibited positive r_SC_ values both within and between classes,, primarily during the visual and delay epochs of the task. During the visual epoch of visually guided saccades (Figure 3A, left), high r_SC_ was observed for neurons within the Visual-Movement class, within the Delay class, and between them. Only modest levels of r_SC_ were observed for other class permutations, with Movement class neurons forming the weakest correlations across classes. Overall, within-class r_SC_ was higher than between-class (p < 0.001, Student’s t-test). A qualitatively similar pattern was observed for r_SC_ during the delay epoch (Figure 3A, middle), but correlated variability during the movement epoch was substantially reduced, both within and between classes (Figure 3A, right). Still, average within-class r_SC_ were higher than between-class (p < 0.001 in both epochs, Student’s t-test). The degree of correlated variability for memory-guided saccades (Figure 3B) followed a similar pattern to that observed for visually guided saccades, but with a conspicuously higher r_SC_ for neurons within the Delay class during the delay epoch, as observed in Figure 2. In addition to measuring r_SC_ between classes within epochs, we measured r_SC_ between epochs. This approach sought to evaluate whether a transfer of information took place over time, but all between-epoch correlations were small and observed for only a limited number of class combinations (Supplementary Figure 3).

Overall, the correlated variability exhibited by SC neurons was predominantly between pairs of neurons within the same functional class and during a given epoch. Pairs from different classes also exhibited significant r_SC_, albeit to a lesser extent.

### r_SC_ in Delay class neurons was modulated by saccade condition and target location

The qualitative difference between visually and memory-guided saccades during the delay epoch motivated us to compare these data directly (Figure 4A). Juxtaposing the saccade conditions revealed that only Delay class neuron pairs displayed levels of r_SC_ that differed between visually guided and memory-guided saccades (p < 0.001, 2-way ANOVA). The relative increase in r_SC_ during memory-guided saccades cannot be due to a difference in firing rate because spike counts were not significantly different between the two saccade conditions, and if anything, were lower for memory-guided saccades (Supplementary Figure 4). Thus, Delay class neuron pairs stood out not only because r_SC_ in this class was uniquely modulated by saccade task condition, but also by virtue of exhibiting the largest level of r_SC_ overall.

**Figure 4:**
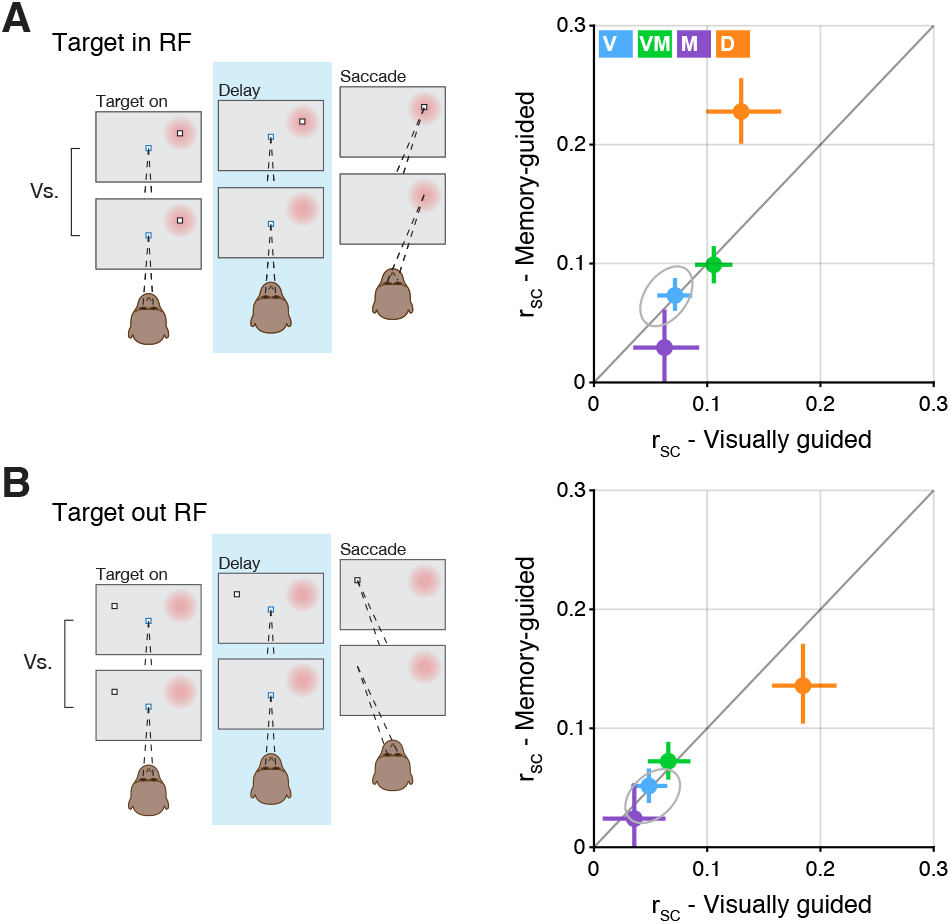
r_SC_ in Delay class neurons was modulated by saccade condition and target location. **(A)** Left: graphic indicting the comparison of visually guided versus memory-guided saccade trials during the delay epoch when the saccade target was presented within the RF of the recorded neurons (RF indicated by the red patch). Right: r_SC_ values in the memory-guided condition plotted against values in the visually guided condition. Error bars indicate bootstrapped 95% confidence intervals on the mean of r_SC_. Gray ellipse reflects bootstrapped 95% confidence intervals on the mean of r_SC_ values for pairs drawn from a class-blind “null” distribution. **(B)** Same format as A, but for when the saccade target was presented outside the RF of the recorded neurons.

The larger r_SC_ for Delay class neurons held true even when target location was outside their RFs (Figure 4B). We found that for “out-RF” trials, during which the saccadic target was presented in the opposite hemifield, SC neurons exhibited a similar pattern to that observed for in-RF trials (Figure 4A), in that Delay class neurons exhibited the highest level of r_SC_ and were uniquely modulated by saccadic condition (p < 0.001, 2-way ANOVA). A key difference between the out-RF and in-RF data, however, is that the sign of the condition-related modulation was inverted: for in-RF trials, Delay class r_SC_ was larger for memory-guided saccades compared to visually guided (Figure 4A), but for out-RF trials, was smaller (Figure 4B). Here too, the difference in r_SC_ cannot be attributed to a difference in firing rate because spike counts were not significantly different between visually and memory-guided saccades (Supplementary Figure 4B).

Thus, the level of r_SC_ in Delay class neurons depended on both saccade condition and target location, consistent with a spatially selective modulatory input related to the difference in task demands between memory versus visually guided saccades.

### No significant r_SC_ detected across hemispheres

The dependence of r_SC_ on target location motivated us to test whether our findings extended to SC neurons across hemispheres. We hypothesized that the relative inversion of Delay class r_SC_ during visually versus memory-guided saccades as a function of target location might indicate inter-hemispheric competition amongst Delay class neurons. On a subset of sessions (n = 6 in one monkey) we recorded from the left and right SC simultaneously (Figure 5A). We used these bilateral neuron pairs to test whether particular functional classes exhibited correlated (or anti-correlated) activity between the two SCs. For example, the presence of inter-hemispheric competition might be expected to produce negative r_SC_ values.

**Figure 5:**
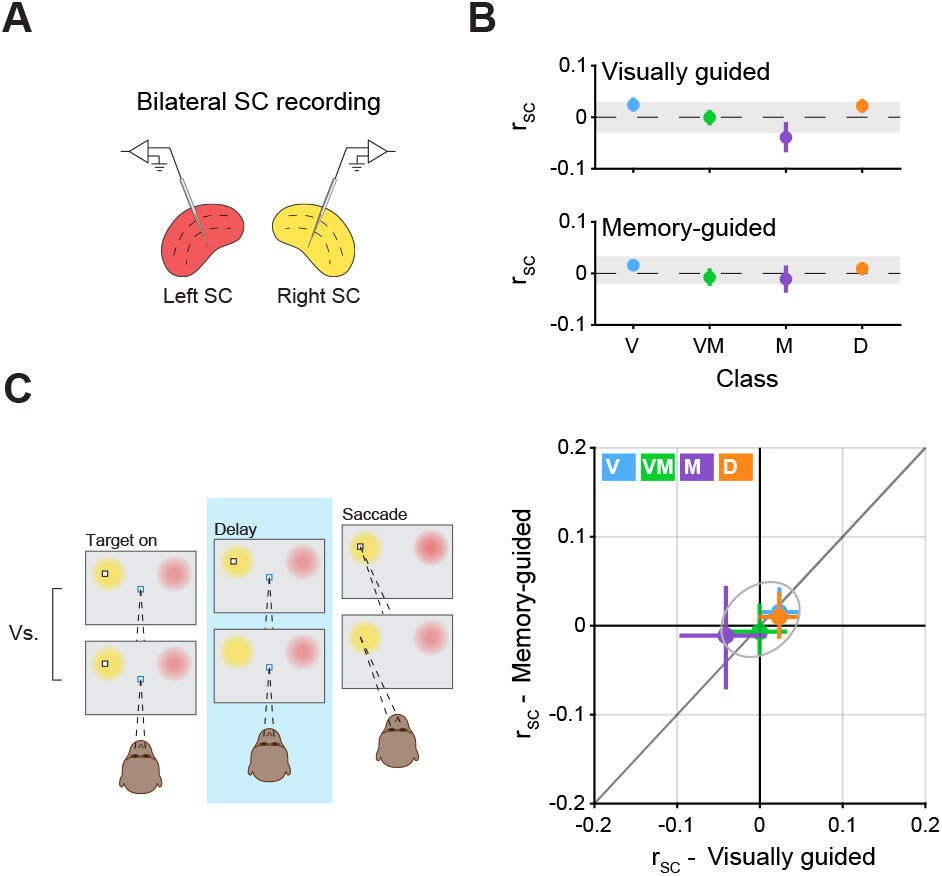
No r_SC_ detected across hemispheres. **(A)** Schematic of bilateral recordings. One multi-channel probe was positioned in each SC, simultaneously recording bilateral pairs of SC neurons. **(B)** Mean r_SC_ values for bilateral pairs in each neuronal class during the delay epoch of visually guided (top) and memory-guided (bottom) saccades. Error bars indicate 1 SEM, bootstrapped. Gray bars indicate bootstrapped 95% confidence intervals on the mean of r_SC_ values for pairs drawn from a class-blind “null” distribution. **(C)** Left: graphic illustrating the comparison of visually guided versus memory-guided saccade trials during the delay epoch when the saccade target was presented in either the right or left set of neuronal RFs, corresponding to either the left or right SC (the graphic shows the target on the left, but trials from both locations were used). Right: r_SC_ values in the memory-guided condition plotted against values in the visually guided condition. Error bars indicate bootstrapped 95% confidence intervals on the mean of r_SC_. Gray ellipse indicates bootstrapped 95% confidence intervals on the mean of r_SC_ values for bilateral pairs drawn from a class-blind “null” distribution.

We found no significant cross-hemisphere r_SC_ for either functional class, regardless of saccade condition (Figure 5B). For each class, no significant difference between the delay epoch r_SC_ and the trial-shuffled (null) distribution (p > 0.05, Student’s t-test) was detected, in either visually or memory-guided saccade trials. Likewise, no significant differences were found between classes in either saccade condition (all p > 0.05, ANOVA). Thus, the correlated variability across bilateral pairs in all classes and saccade conditions was statistically indistinguishable from zero and bounded by values obtained in a power analysis (Methods), indicating that common input onto SC neurons was neither correlated nor anti-correlated, and largely independent across the two hemispheres.

As opposed to r_SC_ within a single SC, where significant differences were observed between classes that depended on saccade condition and target location (Figure 4), no differences between saccade conditions were observed for cross-hemisphere r_SC_, in any functional class (Figure 5C, all p > 0.05, Student’s t-test). This null result is especially important for the Delay class in which a significant difference in r_SC_ was observed between saccade conditions that reversed as a function of target location (Figure 4). Thus, the modulatory effect of saccade condition on Delay class r_SC_ likely operates on the left and right SC independently, with little to no competition across hemispheres.

### Delay class neurons exhibited the longest temporal autocorrelations

Finally, we assessed whether the intrinsic timescale of neuronal activity (Murray et al., 2014) might also differ across the classes of SC neurons. For example, the timescale of processing might be shorter for Movement class neurons that control saccade direction and amplitude at high temporal precision. The intrinsic timescale of Delay class neurons, in contrast, might be expected to be longer, given their role in cognitive functions such as decision-making, selective attention, and working memory (Krauzlis et al., 2013; Basso and May, 2017). We adopted an approach used previously to compute temporal autocorrelations across a hierarchy of brain regions (Murray et al., 2014) and applied it to the different classes of SC. Unlike our previous analyses that focused on either visual, delay or movement epochs of the saccade tasks, this analysis was restricted to a 500ms epoch before target onset during fixation, termed the “foreperiod”. Temporal autocorrelation during this foreperiod can be used to gain insight into the intrinsic timescale of a circuit, independent of task-related factors such as saccade condition or target location.

On average, neurons from each class exhibited a general decay pattern characteristic of a temporal autocorrelation function (Figure 6A). A similar pattern was observed for individual neurons and over all time bins (Supplementary Figure 6A and 6B, respectively). We fit a decaying exponential to the average data to quantify the intrinsic timescale for each functional class. Timescales differed significantly across classes (p < 0.001, Kruskal Wallis), with Movement class neurons exhibiting the shortest intrinsic timescale and Delay class neurons exhibiting the longest, in line with the functional specializations of these neurons. Taken together, results from this analysis of single units complement those obtained by the analysis pairs, whereby both intrinsic timescales and r_SC_ were different across the functional classes of SC, and most pronounced in neurons belonging to the Delay class.

**Figure 6:**
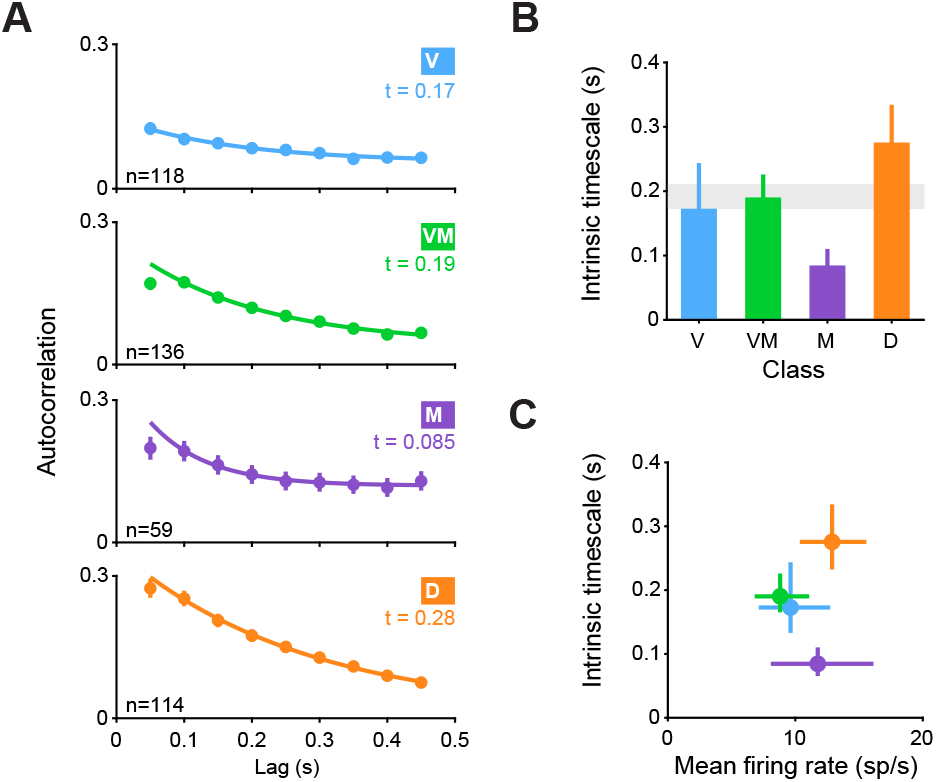
Temporal autocorrelations were longest in D class neurons. **(A)** Spike-count autocorrelations for each functional class of SC neurons. Autocorrelations were computed in 10 non-overlapping, successive 50 ms bins during the foreperiod (during fixation, prior to target onset). Error bars indicate 1 SEM, bootstrapped. Solid line represents the fit of an exponential decay with an offset. Timescale of decay (“t”) is noted on the panel of each class. **(B)** Timescale of decay for each class. Error bars indicate bootstrapped 68.2% confidence intervals on the timescale parameter estimate. Gray bar indicates bootstrapped 68.2% confidence intervals on the timescale parameter estimate computed over all SC neurons irrespective of class. **(C)** Timescale of decay for each class as a function of mean firing rate. Vertical error bars are identical to those in B, horizontal error bars indicate 1 SEM, bootstrapped.

## Discussion

Correlated variability (spike count correlations, r_SC_) is widely reported in the literature and depends on neuronal properties and behavioral factors (Cohen and Kohn, 2011; Kanitscheider et al., 2015; Kohn et al., 2016; Nienborg et al., 2012). In this paper we demonstrated that in addition to these properties, r_SC_ depends on functional class. First, r_SC_ within a functionally identified subpopulation of neurons was strikingly different from the r_SC_ for the overall population of neurons (Figure 2). Second, the amplitudes of r_SC_ varied across different epochs of the task for neurons both within and between classes (Figure 3). Third, Delay class neurons stood out as particularly correlated during the delay epoch of the saccade task, the magnitude of which was modulated by whether the saccade was memory or visually guided, and by whether it was directed into or out of the neuronal RFs (Figure 4). Fourth, despite the dependence of Delay class r_SC_ on saccade direction, no significant r_SC_ (neither positive nor negative) was detected for bilateral pairs of Delay class neurons (or for any other class) (Figure 5). Lastly, Delay class neurons exhibited the largest temporal autocorrelations across all classes (Figure 6).

### Distinct levels of r_SC_ in the SC

Our main finding, that different functional classes in SC exhibit distinct levels of r_SC_ (figure 2), highlights the importance of taking functional class into account when measuring and interpreting r_SC._ The current study investigated primate SC, but microcircuits reflecting distinct functions, processing stages, or associations with feedforward or feedback projections exist in most areas of the brain (Brown and Hestrin, 2009; Burgalossi et al., 2011; Gilbert and Li, 2013; Sharpee, 2014). Thus, while existing measurements of r_SC_ in many brain regions provide important information on the level of coordinated activity over all neuron pairs (Cohen and Kohn, 2011), it is important to consider the possibility that this overall level may obfuscate the actual r_SC_ magnitude distributed over multiple microcircuits within the region. This is especially important in early sensory areas where the measured r_SC_ is taken to inform (and constrain) downstream computations that support behavior, and even more consequential if the distinct level of r_SC_ is in the output neurons.

Why would different classes of SC neurons exhibit distinct levels of r_SC_? Traditionally, the source of r_SC_ was thought to arise from shared noise on the sensory inputs, but recent evidence from sensory cortex has shown that feedback projections play a substantial role in shaping the structure of r_SC_ (Bondy et al., 2018; Cohen and Newsome, 2008; Quinn et al., 2021). Indeed, cortical laminae associated with different circuit positions and functions possess distinct degrees of r_SC_ (Hansen et al., 2012; Smith et al., 2013). Thus, different levels of r_SC_ in a neuronal population correspond to distinct inputs that can be either correlated or decorrelated (Ecker et al., 2010). Our data are consistent with this idea given that different classes of SC neurons reflect distinct functions and processing stages, operate on different timescales (Figure 6), and are associated with different inputs (reviewed in (Wurtz and Albano, 1980; Basso and May, 2017). For example, neurons in the superficial layers of SC (predominantly classified as Visual) receive a preponderance of feed-forward inputs from early processing stages such as retinal ganglion cells, thalamus, and striate cortex, whereas neurons in the intermediate layers (predominately classified as Visual-Movement and Delay) receive inputs from numerous cortical areas, likely involving cortico-subcortical loops and feedback projections. Such a functional architecture is consistent with our finding that changing the saccadic condition from visually guided to memory-guided, which relies on working memory and is thought to engage feedback or recurrent activity (Funahashi et al., 1989; Constantinidis and Wang, 2004; Goldman-Rakic, 1995; Miller et al., 2018), resulted in different levels of r_SC_ in Delay neuron pairs, but not others (Figure 4).

Distinct functional classes have also been identified in FEF and LIP neurons (Bruce and Goldberg, 1985; Gnadt and Andersen, 1988), but a systematic measurement of r_SC_ in each has not been performed. In one related study, distinct levels of r_SC_ were reported in neuronal subpopulations of FEF binned along a visuomotor spectrum (Khanna et al., 2019), consistent with the results reported here. Two important differences, however, are that the Khanna et al. study did not include Delay class neurons as a separate functional class in its analysis and used only memory-guided saccades, thereby precluding an evaluation of whether r_SC_ in some neuronal classes are differentially affected by task manipulations. It is possible that our finding of higher r_SC_ levels in SC Delay class neurons would extend to the analogous classes of neurons in FEF and LIP as well. It is further possible that the higher r_SC_ exhibited within vs. between class (Figure 3) extends to pairs of neurons across areas, such that Delay class neurons in oculomotor areas SC, FEF, and LIP, for example, correlate with one another to a larger extent, but as far as we know, these possibilities have yet to be examined.

### The effect of task on r_SC_ in Delay class neurons

Experimental manipulations aimed at influencing cognitive states have significant effects on r_SC_ in many brain regions (Cohen and Kohn, 2011; Doiron et al., 2016). In the current study, manipulating whether the subject was to perform a visually or memory-guided saccade also influenced r_SC_, but only in Delay class neurons (Figure 4a). Why did Delay class neurons exhibit higher r_SC_ during the delay epoch of memory-guided saccades compared to visually guided? There are two differences between saccade conditions that could contribute to the differences in r_SC_: (1) a target remains in the RF of the SC neurons during visually guided saccades, but not in memory-guided saccades, and (2) the absence of the target during memory-guided saccade trials requires the target position to be maintained in working memory. The presence of the saccade target in visually guided trials represents a potential sensory source of common input that could influence r_SC_ (Churchland et al., 2010; Kohn and Smith, 2005; Smith and Kohn, 2008), but it is unclear why this influence should be found specifically in Delay class neurons, and not in neurons from other classes that process visual information. Instead, it seems more likely that the relative increase of r_SC_ in Delay class neurons during the delay epoch is due to correlated input related to the maintenance of target position in working memory.

What might be the source of such correlated input to Delay class neurons during working memory periods? If correlations were present in the sensory input – that is, due to “noise correlations” as originally defined – then higher r_SC_ in Delay class neurons would indicate that the accuracy of the information they provide as a population about target location would be limited by this shared noise. An alternate view, which we favor, is that the r_SC_ is introduced specifically onto these neurons by other circuits in the brain to meet a behavioral goal, such as maintaining target position in working memory, consistent with recent evidence that r_SC_ are task-dependent (Bondy et al., 2018). Prefrontal and parietal neurons are candidate sources of such correlated input, as these cortices have long been associated with working memory (Goldman-Rakic, 1995; Leavitt et al., 2017; Miller et al., 2018) and exhibit the same type of persistent activity displayed by SC Delay class neurons: activity during the delay period between vision and action, in the absence of a stimulus in the neuronal RF (Wurtz and Goldberg, 1972; Glimcher and Sparks, 1992; McPeek and Keller, 2002; Munoz and Wurtz, 1995). Functional inputs from prefrontal and parietal cortices to SC have been identified previously (Everling and Munoz, 2000; Inoue et al., 2015; Paré and Wurtz, 1997; Sommer and Wurtz, 2000), and inactivation of the frontal eye fields (FEF) has been shown to reduce the number of spikes emitted by neurons located specifically in the intermediate layers of the SC (Peel et al., 2020), which likely correspond to our Delay class neurons. Thus, corticotectal inputs from prefrontal and parietal areas could underlie our observation of high r_SC_ in Delay class neurons that is modulated by the reliance on working memory given the saccade condition.

The direction of r_SC_ modulation by saccade condition depended on whether the saccadic target was in the RF of the neurons under study. When saccades were directed into the RF, r_SC_ was high for memory-guided saccades and low for visually guided (Figure 4a) but when saccades were directed to the opposite hemifield, the reverse was observed (Figure 4b). If input from upstream regions such as prefrontal cortex is the source of increased r_SC_ during memory-guided saccades into the RF, then a similar input must reach the SC on the other side of the brain during out-RF saccades. Such an asymmetry in modulatory input is consistent with inter-hemispheric competition and may produce negative r_SC_ values across the two colliculi, but this was not observed in either class (Figure 5). A push-pull relationship documented in the context of saccades may still be present between certain SC neurons, but this is primarily manifest in spiking output, not modulatory input (Hilgetag et al., 1999; Sprague, 1966). Thus, the modulatory input onto Delay class neurons appears largely independent across the left and right SC. If the source of such input is cortical, this indicates that two independent processes are required in cortex to hold target position in working memory, even though the target appeared on only one side.

### Difference in r_SC_ across classes could not be explained as byproducts of other factors

Measurements of r_SC_ are sensitive to a number of confounding factors such as firing rate and measurement window duration (Cohen and Kohn, 2011). We performed several control analyses to rule out such confounds and found that our main results were unchanged (Supplementary Figure 2, Supplementary Figure 4). Another potential confound in our measurements are small-amplitude fixational eye movements termed microsaccades. Moving the eyes can synchronously modulate the spiking of simultaneously recorded neurons, providing a possible source of pairwise correlations (McFarland et al., 2016). However, movement of the eyes would be expected to primarily affect neurons with visual or movement properties such as neurons from the Visual, Movement, and Visual-Movement classes, and yet these had lower r_SC_ values than neurons from the Delay class. We also considered whether the higher r_SC_ in Delay neurons during memory-guided saccade trials (Figure 4) was due to a larger proportion of microsaccades or other behavior not specifically controlled for during these trials compared to visually guided. But again, this explanation would seem to imply increases in r_SC_ in all classes during memory-guided saccade trials, not just Delay class neurons. And would imply similar effects on r_SC_ for both in-RF and out-RF trials, and in both colliculi, in contradiction to our findings (Figure 4; Figure 5). The same argument can be used to rule out global factors related to arousal, as mechanisms related to arousal would be expected to influence both in-RF and out-RF trial types, as well as both colliculi. Thus, while the occurrence of microsaccades and non-specific arousal signals may affect overall levels of r_SC_, it is unlikely that these could account for the differences across classes and tasks.

Of course, it is important to note that the distinction between neuronal classes is not clear cut, and the response properties of a neuron lie on a spectrum (Gandhi and Katnani, 2011). In fact, a recent study (Khanna et al., 2019) took this point into consideration and binned neuronal function in FEF on a visual-movement spectrum to avoid categorical classifications. However, reducing the space to one axis or dimension cannot capture unique functional signatures such as persistent activity, which can manifest in neurons regardless to whether they are predominantly visual, movement, or in between. Thus, the discrete classification approach taken here, while coarse, facilitated the identification of specific neuronal classes that exhibited unique pairwise responses during certain epochs of visually and memory-guided saccades.

## Summary

As recording techniques advance and larger pools of neurons are recorded simultaneously, considering r_SC_ as a metric of functional connectivity has become standard practice. Gaining better insight into what constitutes a population of neurons or whether a population is composed of several sub-populations is important for accurately measuring r_SC_ in a brain area and informing population coding. Here we found that neurons identified by functional class exhibited very different levels of r_SC_, presumably because each sub-population of neurons occupies a distinct circuit niche within the SC network with distinct modulatory inputs. Our findings highlight the importance of taking neuronal class into account when measuring and interpreting the significance of correlated activity across different tasks and conditions.

## Acknowledgements

We thank Nick Nichols, Daniel Yochelson, Denise Parker and Amber Lopez for technical support. We are grateful to Hendrikje Nienborg and Bruce Cumming for providing feedback on an earlier version of the manuscript, and to Christian Quaia, Lupeng Wang, Kerry McAlonan and Kara Cover for helpful discussions. This work was supported by the National Eye Institute Intramural Research Program at the National Institutes of Health (ZIA EY000511).

## Author contributions

L.N.K, G.Y., J.P.H. and R.J.K. designed the experiments. L.N.K and G.Y. conducted the experiments. L.N.K analyzed the data. L.N.K and R.J.K drafted the manuscript. All authors edited and revised the manuscript.

## Methods

### Animals

Two adult male rhesus monkeys (*Macaca mulatta*) weighing 9-12 kg were used in the study. All experimental protocols were approved by the National Eye Institute Animal Care and Use Committee and all procedures were performed in accordance with the United States Public Health Service policy on the humane care and use of laboratory animals. A plastic headpost and recording chamber had been previously implanted granting electrophysiological access to the superior colliculus (SC).

### Experimental apparatus

Animals were seated and head-fixed in a primate chair (Crist instrument Inc) inside a darkened booth facing a VIEWPixx display (VPixx Technologies) that was controlled by a mid-2010 Mac Pro (Apple Inc) running MATLAB (The Mathworks) with the Psychophysics Toolbox extensions (Brainard, 1997). Eye position was recorded using an EyeLink 1000 infrared eye-tracking system (SR Research Ltd.) and monitored online for gaze-contingent progression through the task. Experiments were controlled using a modified version of PLDAPS (Eastman and Huk, 2012).

### Guided saccade task

Animals performed a saccade task that included both visually guided and memory-guided saccade trials (figure 1b). In either version of the task, a trial began with the appearance of a 0.25° wide fixation square (48 cd/m^2^) on a gray background (28.5 cd/m^2^). Fixation had to be maintained within a 2° wide square window (invisible to the monkey). At 0.5 - 0.7 s following fixation acquisition, a saccade target appeared in one of four possible locations and either stayed on (visually guided condition) or was extinguished following 0.2 s (memory-guided condition). Monkeys were required to maintain fixation during target presentation up until the disappearance of the fixation square (the “go signal”, 1 - 2 s following target onset), whereupon a saccade was to be executed towards the target within 0.1 – 0.5 seconds, land within a 3° wide window around the target (invisible to the monkey) and maintained within the window for a duration of 0.5 – 0.7 seconds. In the memory-guided condition, the target was reillumined following entrance into the target window for a duration of 0.2 seconds. Successful completion of a trial resulted in a juice reward while any deviation aborted the trial and trial identity (i.e. target location and condition) was reshuffled into the block of trials. The precise timings of events on every trial were determined by random draws from a uniform distribution within the prescribed ranges.

Each block consisted of 24 trials distributed across four target locations (one pair of targets in each hemifield). In each hemifield, one target was positioned to maximally overlap with the response fields (RF) of neurons recorded in the contralateral SC, and the other at a 90° angle away from the first either above or below the horizontal median, at a similar radius. In sessions where only one SC was recorded from, targets on the ipsilateral hemifield were positioned diametrically opposite to those in the contralateral. Trials were distributed at a proportion of 2:1 for each pair, where targets placed in neuronal RFs were used more frequently. Trial condition was distributed equally (i.e., 50% visually guided and 50% memory-guided) for each target location. Overall, a typical session consisted of 20 blocks and netted 516 ± 126 trials (mean ± std).

### Electrophysiological recordings

Electrophysiological signals and analog input (e.g., eye position, joystick position) were acquired by an Omniplex-D system (Plexon Inc.). Neuronal activity in SC was recorded using 32-channel Plexon v-probes with 50µ inter-channel spacing (Plexon Inc.) from either one of the SCs or both simultaneously, with one probe in each SC. Probes were advanced to their target depth in the intermediate and deep layers of the SC using a motorized microdrive (NAN Instruments). Target depth was guided by previous mapping and verified by measuring neuronal responses online. Activity on each channel of the probe was thresholded (µ - 3σ) and used to map the spatial RF by having the animal perform visually guided saccades to targets presented at locations drawn randomly from an XY grid (5° spacing from −25° to 25° on the X-axis, and −15° to 15° on the Y) as well as locations set manually by the experimenter to more precisely measure the spatial extent of the RF. A saccade target was then positioned in the center of the RFs and used to collect several memory-guided saccades (typically ∼20) sufficient to ascertain the probe’s location within the SC: visual responses were associated with superficial SC layers; saccade-related activity with deep. Probe position was then adjusted to maximize neuronal yield from intermediate and deep SC layers, and left to settle (∼1 hour) to allow the tissue to stabilize before beginning the experimental session.

### Electrophysiological analysis

Continuous spike-channel data collected during the experimental session were sorted offline with Kilosort2 (Pachitariu et al., 2016) and manually curated by a human expert using Phy2 to ensure that all sorted units have plausible inter spike interval distributions and waveform shapes consistent with action potentials. Neurons with low (<1.5) signal-to-noise ratio (Kelly et al., 2007) were excluded. Overall, 366 neurons were recorded from 16 sessions collected in Monkey #1, and 542 neurons from 29 sessions in Monkey #2. Offline, we determined the “in-RF” target for each neuron as the target that elicited the largest visual (+50 to +200 ms relative to target onset) or movement (25 ms before saccade onset to saccade end) responses for each neuron. If the visual or movement related responses were not significantly larger than baseline (−75 to +25 relative to target onset) using an ANOVA (alpha = 0.05, corrected), the neuron was not associated with an in-RF target. Only neurons associated with an in-RF target were used for subsequent analysis. We further excluded low firing neurons (<0.1 spikes/s averaged over all targets and time) as rSC estimates for low firing neurons tends to be poor (Cohen and Kohn, 2011). Increasing this exclusion criteria to 1 or 5 spikes / s did not change the main results. Overall, 751 of 908 neurons were used for subsequent analysis. Results did not differ across monkeys and were therefore combined to increase statistical power.

For the visualization of mean firing rated over time relative to key events in the task (Figure 1E), normalized spike counts were binned into non-overlapping 20 ms bins. Each neuron’s data was z-scored normalized by subtracting the mean and dividing its binned spike counts by the standard deviation of that neuron’s counts across trials and conditions.

### Neuronal classification

Each neuron was assigned a functional class following established criteria (Basso and Wurtz, 1998; Herman and Krauzlis, 2017; McPeek and Keller, 2002). Briefly, spike counts during four epochs in the memory-guided saccades to the in-RF target were used to determine neuronal class: baseline epoch (−75 to +25ms relative to target onset); visual epoch (+50 to +200 ms relative to target onset); delay epoch (−150 to +50 relative to fixation offset); movement epoch (−25 ms from saccade onset to saccade offset). Saccade onsets and offsets were calculated offline using a velocity threshold and verified by inspection. A one-way Kruskal-Wallis nonparametric ANOVA was used on spike counts in the four epochs to determine whether a neuron possessed visual-, delay-, or movement-related responses. Such an analysis netted seven possible classes: visual (*v*), visual-delay (*vd*), visual-movement (*vm*), visual-delay-movement (*vdm*), delay (*d*), delay-movement (*dm*), and movement (*m*). This classification was used to construct four operational classes that correspond to past classifications of SC neurons: a “Visual” class, consisting of *v* neurons (29% of our population); a “Visual-Movement” class, consisting of *vm* neurons (27%); a “Movement” class, consisting of *m* neurons (15%); and a “Delay” class, consisting of all neurons that exhibited delay-related activity: *vd* (5%); *vdm* (9%); *d* (1%); and *dm* (2%). The Delay class is similar to “prelude” (or “build up”) neurons (Basso and Wurtz, 1998; Dorris et al., 1997; Glimcher and Sparks, 1992; Herman and Krauzlis, 2017; McPeek and Keller, 2002) as it includes similar groups of neurons (the *vdm* and *dm*), but is distinct in that the Delay class used here also includes *vd* and *d* neurons. Whether or not our Delay class included all delay-exhibiting neurons (*vd, vdm, d* and *dm*) or only a subset corresponding to the classically defined “prelude neurons” (*vdm* and *dm*) did not change any of our main results (see “Delay class subset” in Supplementary figure 2a). Overall, our neuronal dataset includes 751 neurons, of which 220 were classified as Visual, 203 as Visual-movement, 114 as Movement, and 135 as Delay (Figure 1D). 79 neurons exhibited no distinct activity in either epoch of the task and were not used for further analysis.

### Spike count correlation (r_sc_) measurements

Spike count correlations between pairs of simultaneously recorded neurons (r_sc_) were defined as the Pearson’s correlation coefficient of spike counts during repeated instances of the same task conditions, calculated as:

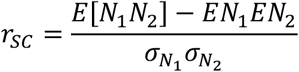

where *N*_1_ and *N*_2_ are the spike counts for neurons 1 and 2 (respectively), *E* is the expected value of the counts, and *σ* is the standard deviation. We followed standard practice to avoid contamination in estimations of r_sc_ values by removing trials with response outliers (>3 SDs difference from the mean) (Kohn and Smith, 2005; Zohary et al., 1994). All statistical evaluations of r_sc_ values within or between classes were performed following the Fisher r-to-z transformation to approximates normality and stabilize variance in the data (Anderson, 1984; Kohn and Smith, 2005). Untransformed r_sc_ values were only presented when visualizing individual pairs (supplementary figured 1 and 2b).

Only neurons with overlapping RFs were included in our analysis of pairwise correlations, netting 5339 pairs overall. For pairs of neurons within the same class, we obtained 627, 590, 144, and 280 pairs for the Visual, Visual-movement, Movement, and Delay classes, respectively. r_sc_ values were computed at specific epochs with a duration of at least 150 ms (see shaded regions in figure 2a) and over time (time courses in figure 2a-d), using a 200 ms sliding window with 50 ms increments.

### Spike count autocorrelation measurements

Measurements of spike count autocorrelation (figure 6 and supplementary figure 5) were performed following a procedure described previously (Murray et al., 2014). Analysis was performed on the foreperiod, a 500ms epoch during the fixation period that preceded target onset. In this epoch, animals were in a controlled state of preparedness with no variation in sensorimotor features of the task across trials. The foreperiod was divided into 10 separate, successive 50 ms bins and the Pearson’s correlation coefficient was computed between each *i*-th and *j-*th bin. For a single neuron or the average over a population, this procedure produces an autocorrelation matrix (Supplementary figure 5a). Decay of the temporal autocorrelation was computed by fitting a decaying exponential with an offset as a function of the time lag *k*Δ between bins (*k* = |*i* − *j*|):

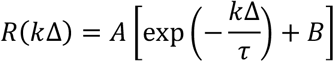

where *τ* is the time constant reflecting the intrinsic timescale of decay, and *B* is the offset. Some neurons exhibited a lower autocorrelation in the first time lag (50 ms) compared to subsequent lags, consistent with refractoriness. This has been noted previously (Murray et al., 2014) and overcome by fitting the exponential decay to data starting at the largest reduction in autocorrelation between two consecutive time bins and onwards. Temporal autocorrelations in Figure 6 were averaged over neurons and lags. Temporal autocorrelations averaged over neurons but not lags, and over lags but not neurons, are presented in Supplementary figures 5a and 5b, respectively.

### Pairwise signal correlation measurement

Measurements of signal correlations across pairs of simultaneously recorded neurons (Supplementary figure 2B) were performed on data obtained during the mapping of spatial RFs described above. In the task, visually guided saccades were executed towards many locations in space with variable proximity to the RF center of the neurons under study, thereby introducing variations in signal. Signal correlation was computed by calculating the Pearson correlation between responses of a pair of neurons during visually guided saccades (from target onset to saccade offset) to the range of targets presented.

### Statistical analyses

To determine whether different classes of SC neurons exhibited different levels of r_sc_ standard tests such as Student’s t and ANOVA were used (Bonferroni corrected for multiple comparisons). This choice was made because r_sc_ values used for statistical testing underwent a Fisher r-to-z transformation, which approximated normality and stabilized variance in the data (Anderson, 1984; Kohn and Smith, 2005). Using non-parametric tests instead (e.g. Wilcoxon Rank Sum or Kruskal-Wallis analysis of variance) did not change the results. Confidence intervals and SEMs were estimated using standard a bootstrap procedure with 10,000 random draws with replacement. To determine the statistical significance of a distribution of r_sc_ values the distribution was compared to a matched distribution in which trial identity was shuffled (Supplementary figure 1A). For class-blind “null” distributions (gray band or ellipse in Figure 2G, H, Figure 4A, B, Figure 5B, C), trial identity was not shuffled, but pairs were randomly sampled from the full dataset of 5339 neuronal pairs, irrespective of functional class.

To determine whether the reported differences in r_sc_ values across classes were artifacts of changes in firing rate or due to shorter distances between neurons, a mean-matching procedure (Churchland et al., 2010; Ruff and Cohen, 2014) was implemented (Supplementary Figure 2A). Briefly, firing rates (or distances) of neuronal pairs were binned into 12 equally sized bins across all classes of neurons. From each bin, we randomly sampled (without replacement) a number of values that was determined by the bin with fewest data points, thereby creating sub-distributions with equal values across bins, matching the distribution of firing rates (or distances). This process was repeated 10,000 times to obtain the averages and SEMs presented in Supplementary Figure 2A. Because Movement class neurons fired very sparsely during the delay epoch, their firing rate distribution did not overlap with the distributions of the other classes to a large enough extent, reducing the statistical power substantially. We therefore excluded the Movement class from the mean-matching procedure for firing rate (but not distance).

### Power analysis

In six sessions, we recorded from the SC bilaterally. Overall, 738 pairs of cross-hemisphere neurons were obtained, netting 82, 32, 9 and 52 bilateral pairs from the Visual, Visual-Movement, Movement and Delay functional classes, respectively. In our limited dataset, no correlations were detected for either functional class (Figure 5). To determine the smallest absolute value of r_SC_ that our analysis method could detect with a confidence of 95%, we performed a bootstrapped power analysis. For each individual functional class, we simulated distributions of r_sc_ values with varying means (from −0.2 to 0.2, 0.005 increments) and a SD determined by the SD of the class-blind distribution of bilateral pairs (0.09). We randomly drew a number of samples equal to the number of pairs obtained in our recordings for that class and tested whether the mean r_sc_ was significantly different from zero (Student’s t-test). We repeated this process 10,000 times to identify the mean r_sc_ value at which we could detect a statistical difference in 95% of cases. This approach determined the highest possible value of r_sc_ that could exist in our data and go undetected, with 95% confidence. For the Visual, Visual-Movement, Movement and Delay classes, these values were 0.05, 0.09, 0.17 and 0.07 for visually guided saccades, and 0.05, 0.08, 0.16 and 0.06 for memory-guided saccades, respectively

