## Supplemental Figures for "Correlated variability in primate superior colliculus depends on functional class"

#### Supplementary figure 1: $r_{SC}$ distributions over neuronal classes

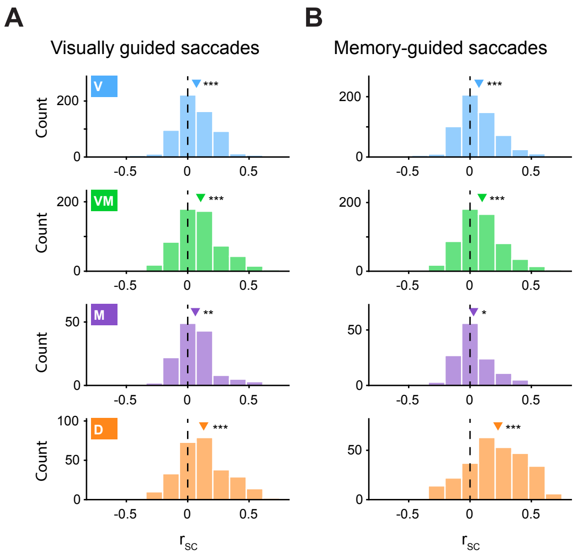


Distributions of $r_{SC}$ values within each neuronal class during the delay epoch of visually- and memory-guided saccades. These data were used to compute the mean $r_{SC}$ and SEM displayed in Figure 2 G-H. **(A)** For visually guided saccades, distribution means (indicated by triangles on figure panels) ± s.d. for neuron pairs in the four classes were: 0.07 ± 0.18; 0.11 ± 0.2; 0.06 ± 0.18; and 0.13 ± 0.23. All four distributions were significantly different from a matched trial-shuffled (null) distribution (Student's t-test, asterisks on panel denote levels of statistical significance: *** $p < 0.001$; ** $p < 0.01$; * $p < 0.05$) **(B)** For memory-guided saccades, distribution means ± s.d. were: 0.07 ± 0.19; 0.10 ± 0.2; 0.03 ± 0.17; and 0.23 ± 0.27. All four distributions were significantly different from a matched trial-shuffled (null) distribution (same format as A).

#### Supplementary figure 2: Dependence of $r_{SC}$ on functional class was not due firing rate, distance or signal correlations

**(A)** We evaluated whether the variation in $r_{SC}$ values across functional class presented in Figure 2G,H was influenced by firing rate, inter-pair distance, or details of functional classification method. We recomputed $r_{SC}$ values during the delay epoch of either visually- (left) or memory-guided saccades (right) for three analysis variations: mean-matched firing rate; mean-matched distance; and Delay class subset. For mean-matched firing rate, the firing rates of each pair of neurons were matched across classes (excluding Movement class, see Methods) according to a standard mean-matching method (Methods). For mean-matched distance, the distance between each pair of neurons was matched across neuronal classes following the same mean-matching method. The location of each neuron was estimated as the channel on which its waveform amplitude was largest, and the distance of a pair was the distance between the two channels. For the Delay class subset analysis, inclusion criteria for what constituted the Delay class differed to include only a subset of neurons within that class based on a different method for neuronal classification (See Methods for further details. Inclusion criteria for all other classes remained unchanged). In all variations of analysis method (firing rate, distance, or Delay class inclusion criteria), the significant difference across classes reported in Figure 2G, H was still observed, both for visually guided saccades ($p < 0.01$, $p < 0.05$, and $p < 0.001$, for the three analysis variations, respectively, ANOVA), and for memory-guided saccades ($p < 0.001$, $p < 0.05$, and $p < 0.001$, respectively, ANOVA).

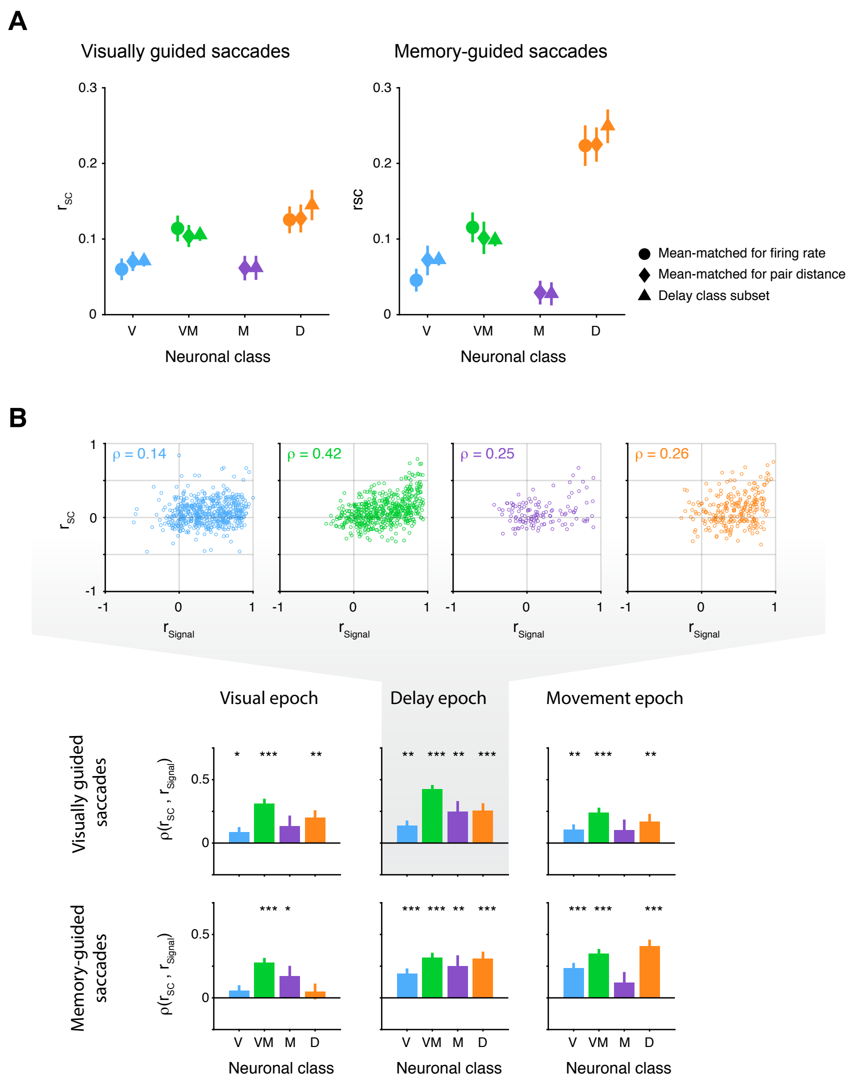


**(B)** We evaluated whether the variation in $r_{SC}$ values across functional class presented in Figure 2G,H was influenced by variations in the relationship between signal correlation ($r_{Signal}$) and $r_{SC}$. Top: the relationship between $r_{Signal}$ and $r_{SC}$ for each functional class. Pearson correlation coefficients (***ρ***) between the two measurements are noted on the panels. Bottom: summary of the correlation coefficients for each class during the visual, delay and movement epochs (columns) and saccade conditions (rows). Asterisks above bars denote the statistical significance of the Pearson correlation: *** $p < 0.001$; ** $p < 0.01$; * $p < 0.05$.

### Supplementary Figure 3: $r_{SC}$ across epochs were present in only a subset of neuronal classes

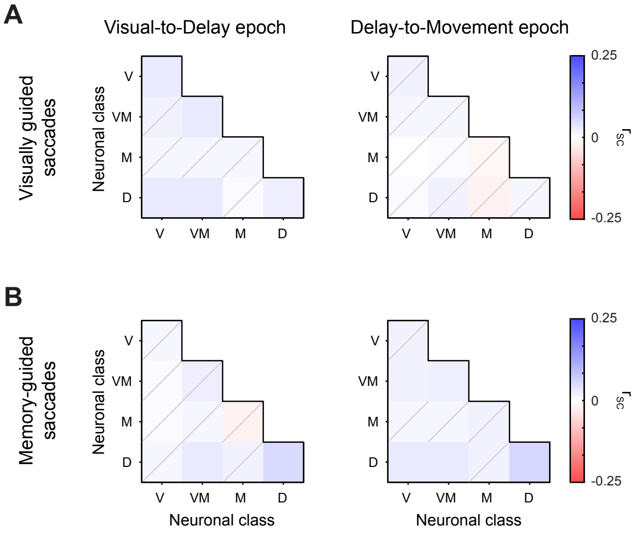


**(A-B)** Heatmaps indicating the mean $r_{SC}$ of neuron pairs between functional classes for visually guided (A) and memory-guided (B) saccades, across epochs. Heatmaps on the left indicate the $r_{SC}$ measured in neuron pairs between the visual and delay epochs, heatmaps on the right indicate the $r_{SC}$ measured in pairs between the delay and movement epochs. Elements on the diagonal of the heatmap indicate within-class $r_{SC}$ values, off diagonal elements indicate between-class $r_{SC}$. Elements for which the measured $r_{SC}$ was not statistically significantly different from their trials-shuffled (null) distribution are indicated by a gray diagonal on the element ($p > 0.05$, Student's t test, Bonferroni corrected).

### Supplementary Figure 4: The difference in $r_{SC}$ between visually and memory-guided saccades did not depend on firing rate.

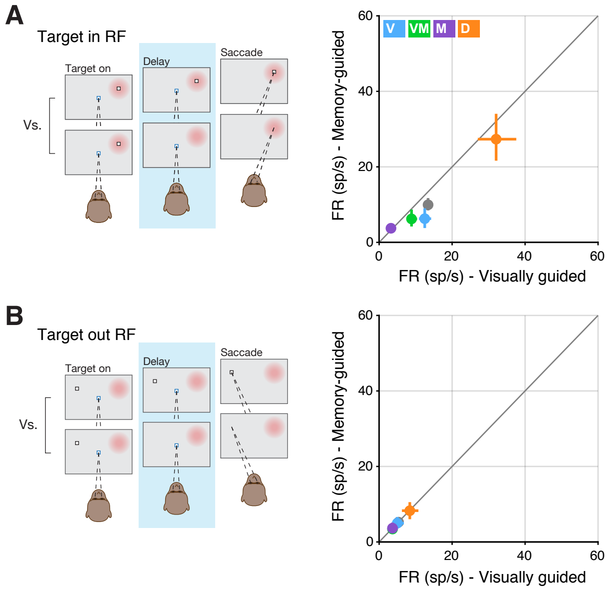


**(A)** Left: graphic indicting the comparison of visually guided versus memory-guided saccade trials during the delay epoch, when the saccade target was presented in the neuronal RF (red patch). Right: mean firing rate (FR) values in the memory-guided saccade condition plotted against FR values in the visually guided condition. Error bars indicate bootstrapped 95% confidence intervals on the mean FR. **(B)** Same format as A, but when the saccade target was presented outside of the neuronal RF.

**Supplementary Figure 5: Spike-count autocorrelations unwrapped in time and for single neurons, across functional class.**

**(A)** Normalized spike-count autocorrelation matrices are presented for each functional class of SC neurons. Matrix elements show the mean correlation of the spike count in each time bin with the spike count in every other bin, averaged across neurons. **(B)** Spike-count autocorrelations are presented for individual neurons within each functional class. Colored markers indicate the population mean (identical to figure 6A). Error bars indicate 1 SEM, bootstrapped. Solid line represents the fit of an exponential decay with an offset (see Methods).

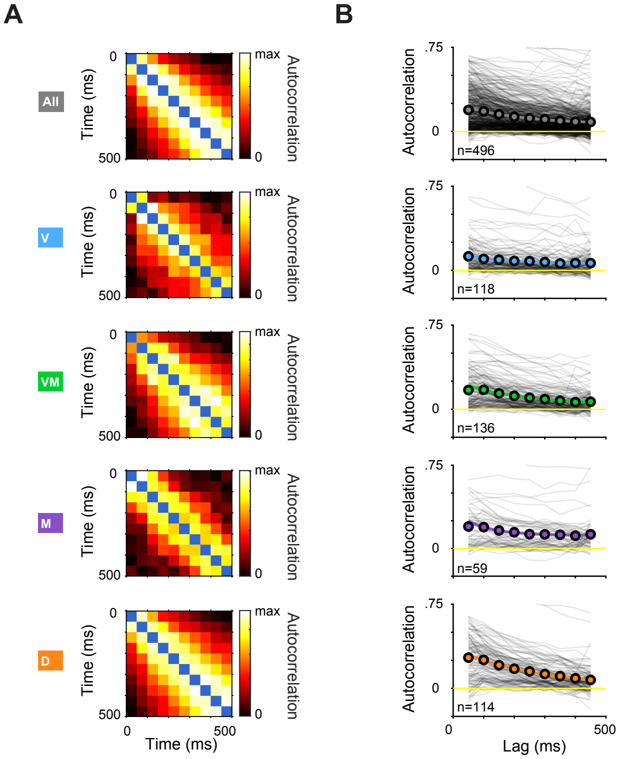
